# PP2A-B55α controls keratinocyte adhesion through dephosphorylation of the Desmoplakin C-terminus

**DOI:** 10.1101/2022.10.19.512916

**Authors:** Abbey L Perl, Jennifer L Koetsier, Kathleen J Green

## Abstract

Critical for the maintenance of epidermal integrity and function are attachments between intermediate filaments (IF) and intercellular junctions called desmosomes. The desmosomal cytoplasmic plaque protein desmoplakin (DP) is essential for anchoring IF to the junction. DP-IF interactions are regulated by a phospho-regulatory motif within the DP C-terminus controlling keratinocyte intercellular adhesion. Here we identify the protein phosphatase 2A (PP2A)-B55α holoenzyme as the major serine/threonine phosphatase regulating DP’s C-terminus and consequent intercellular adhesion. Using a combination of chemical and genetic approaches, we show that the PP2A-B55α holoenzyme interacts with DP at intercellular membranes in 2D- and 3D- epidermal models and human skin samples. Our experiments demonstrate that PP2A-B55α regulates the phosphorylation status of junctional DP and is required for maintaining strong desmosome-mediated intercellular adhesion. These data identify PP2A-B55α as part of a regulatory module capable of tuning intercellular adhesion strength and a candidate disease target in desmosome-related disorders of the skin and heart.

## INTRODUCTION

Cytoskeletal-associated intercellular adhesion junctions are essential for maintaining the stability of multi-cellular tissues and providing cells with the structural integrity to withstand the changing mechanical environment of the tissue. This is particularly important in tissues experiencing high external or internal levels of mechanical stress such as the stratified epidermis or heart. Particularly important for these tissues are desmosomes, intercellular junctions that integrate chemical and mechanical stimuli to allow for the dynamic regulation of the cortical cytoskeleton^1,2^.

The desmosome comprises transmembrane cadherins from two families, desmogleins (Dsg) and desmocollins (Dsc), two plaque armadillo proteins, plakophilin (Pkp) and plakoglobin (PG), and an intercellular filament (IF) cytoskeletal linker protein, desmoplakin (DP). By tethering the IF cytoskeleton to the plasma membrane, DP strengthens adhesion and distributes forces throughout tissues^1,2^. Unlike the Dsg, Dsc, and Pkp families, which are regulated by isoform expression at specific differentiated layers in the stratified epidermis, DP is the only desmosome plakin protein and is therefore ubiquitously expressed in all desmosome-forming cells^3^. The essential nature of DP is best highlighted by the embryonic lethal phenotype of DP null mice, and severe defects exhibited in high-tension tissues including the skin and heart of tetraploid rescue and epidermal specific knockout mice^4-6^. In addition, mutations in the DSP gene result in a range of disorders from lethal acantholytic epidermolysis bullosa (LAEB) to striate palmar plantar keratoderma (SPPK) as well as arrhythmogenic cardiomyopathy (AC) and cardiocutaneous Carvajal syndrome^7-13^.

Notably, DP is regulated in part by the processive phosphorylation of a 68-residue glycine-serine-arginine repeat at its C-terminus directly downstream of the DP-IF binding site (Figure 1A)^14,15^. Previous work from our group and others have characterized the hypo-phosphorylated form of DP as having increased IF binding affinity that impacts desmosome formation, dynamics, and function^14,16-18^. Specifically, expression of constitutively hypo-phosphorylated DP mutants increased adhesion strength and tissue stiffness, resulting in epidermal cell sheets able to withstand higher mechanical stress^17-19^. The phosphorylation of DP’s phospho-regulatory motif was previously identified to be triggered by the coordinated activity of the glycogen synthase kinase 3 (GSK3) and the protein arginine methyltransferase 1 (PRMT-1)^14^. Despite the importance of the hypo-phosphorylated form of DP in strengthening IF binding, the phosphatase responsible for dephosphorylating DP remained unknown.

**Figure 1:**
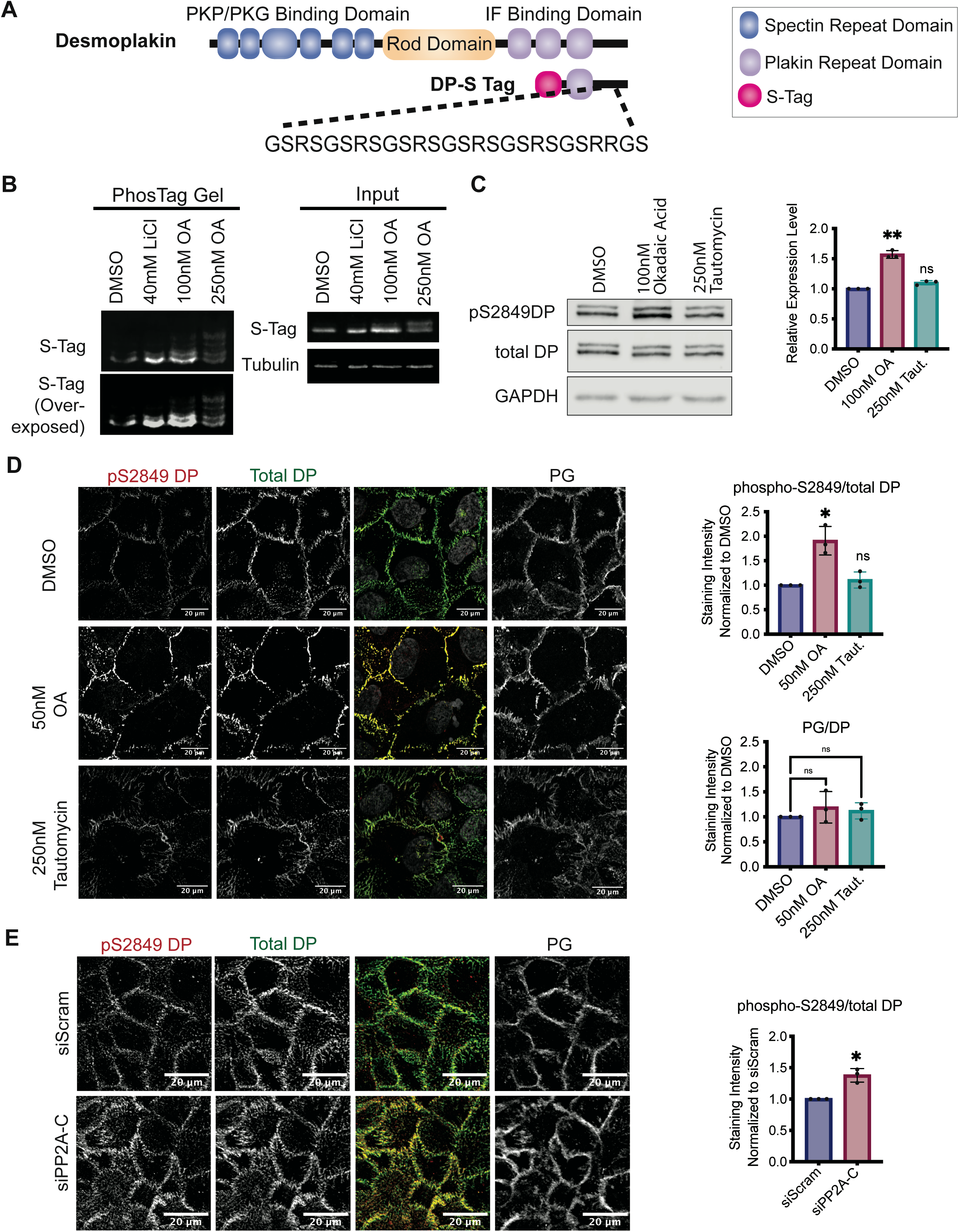
PP2A is a phosphatase for the C-terminal domain of DP. [A] Schematic of DP structural domains and the DP-S-Tag construct containing only residues 2628-2871. Highlighted below is the “GSR” repeat phospho-regulatory motif downstream of the IF binding site capable of regulating DP-IF interactions. [B] A 12% acrylamide gel with (left) and without (right) Phos-Tag molecule capable of separating protein by its number of phosphate groups. Protein lysates are from SCC9 cells transfected with the DP-S Tag C-terminus and treated with either LiCl or OA for 3 hours. [C-E] SCC9s were treated with inhibitors preferential for either PP2A (50 and 100nM OA) or PP1 (250nM Tautomycin) for 3 hours. Expression levels of S2849 phosphorylated and total DP were analyzed in total cell lysates [C] and by immunofluorescence staining [D]. [E] Staining intensity from [C] was quantified at the membrane using PG stain as a mask. Statistical analyses were performed on normalized data using a One sample t comparison to 1. * < 0.05; ** < .01.

This study identifies the protein phosphatase 2A (PP2A)-B55α holoenzyme as the phosphatase responsible for de-phosphorylating DP’s C-terminus. Using a combination of chemical and genetic approaches we show that inhibition of the PP2A-B55α holoenzyme induces an increase in DP phosphorylation specifically at sites of intercellular junctions. Additionally, we find that the PP2A regulatory subunit B55α is associated with DP, particularly at intercellular membranes where both DP and B55α localization is dependent on the other’s expression. Lastly, keratinocyte intercellular adhesion assays showed that loss of the PP2A-B55α holoenzyme diminishes intercellular adhesion strength. Collectively, these data reveal the existence of a previously unrecognized regulatory module capable of fine-tuning desmosome-dependent adhesion. We propose that this module provides a mechanism for rapid adaptations necessary for maintaining a functional barrier in the dynamic mechanical environment of the epidermis.

## RESULTS

### PP2A regulates the phosphorylation of DP’s C-terminus

The importance of hypo-phosphorylated DP for stabilizing the DP-IF interaction to strengthen desmosome-mediated adhesion suggests the existence of a phosphatase that dephosphorylates DP in a regulated matter. To determine if serine/threonine protein phosphatases are involved in this process, keratinocytes were treated with the pan-serine/threonine phosphatase inhibitor Okadaic Acid (OA). To afford comparisons with previous data identifying GSK3 and PRMT-1 as regulating the phosphorylation of DP’s C-terminus, we initially chose to use the squamous cell carcinoma cell line SCC9. SCC9 cells were transfected with a 32 kDa truncated form of DP (DP-S-Tag, Figure 1A) containing only it’s C-terminal domain including residues 2628-2871^14^. DP-S-Tag expressing cells were treated with 100nM OA, 250nM OA, or DMSO vehicle control for 3 hours (Figure 1B). As an internal control, cells were treated with 40mM of lithium chloride (LiCl), a GSK3β inhibitor known to induce a hypo-phosphorylated DP C-terminus^14^. Cell lysates were collected and run on gels containing a Phos-tag peptide capable of separating proteins based on the number of phosphate groups present^20^. The Phos-tag gel showed that LiCl treatment induced an increase of hypo-phosphorylated DP, while both 100nM and 250nM OA treatment increased the level of phosphorylated DP as evident by an accumulation of upper DP-S-Tag bands (Figure 1B).

To identify the phosphatase involved in dephosphorylating DP’s C-terminus, we employed chemical inhibitors targeting the two major serine/threonine protein phosphatases that together account for roughly 90% of all serine/threonine phosphatase activity in keratinocytes, PP2A and PP1. While phosphatase inhibitors exhibit cross-reactivity in their substrates, their preferential selectivity allows for differential inhibition of PP2A and PP1. OA preferentially inhibits PP2A (IC50=∼0.1nM) with an affinity 10x greater than its next major target PP1, (IC50=∼15nM) while Tautomycin preferentially inhibits PP1 (PP1 IC50=∼0.2nM, PP2A IC50=∼1nM)^21^. Therefore, SCC9 cells were treated with either 100nM OA, 250nM Tautomycin, or DMSO vehicle control for 3 hours. A phospho-antibody specific to the S2849 site on DP’s C-terminus was used as a readout for DP C-terminus phosphorylation. OA, but not Tautomycin, treatment increased the levels of phosphorylated DP in total cell lysates (Figure 1C). To determine if PP2A inhibition affects DP phosphorylation at cell-cell junctions, immunofluorescence staining of phosphorylated and total DP in SCC9s treated with drug for 3 hours was performed. As the higher dose of 100nM OA detrimentally affected cell morphology and cell contacts, a lower dose, 50nM OA, that had no visible toxicity or morphological affects was used alongside 250nM Tautomycin. Even with 50nM OA the amount of phosphorylated DP at cell-cell membranes increased almost 2-fold, while 250nM Tautomycin had no detectable impact on the level of phosphorylated DP at the cell membrane (Figure 1D). Moreover, there was no detectable change in the total amount of membrane-associated DP, suggesting changes detected were specific to DP phosphorylation. This is consistent with data generated in the spontaneously immortalized, non-transformed keratinocyte cell line HaCaT (Supplemental Figure 1A) as well as in primary neonatal human epidermal keratinocyte (NHEK) cells (Supplemental Figure 1B). To further confirm the effects on DP’s phosphorylation was specific to PP2A activity, HaCaT cells were transfected with siRNA targeting PP2A’s catalytic subunit (PP2A-C) or a scramble control sequence and grown at confluency for 2 days. Consistent with OA treatment, knockdown of PP2A-C increased the level of phosphorylated DP at the membrane (Figure 1E). Based on these initial data suggesting PP2A, and not PP1, is responsible for regulating DP’s C-terminus, we focused on further elucidating the PP2A-DP relationship.

### B55a exists in complex with Desmoplakin

PP2A is a heterotrimeric protein comprising a scaffolding subunit (A), a catalytic subunit (C), and a regulatory subunit (B). To form the holoenzyme, the A and C subunits bind one of 23 different B subunit isoforms which are responsible for binding the substrates^22^. To better understand how PP2A regulates DP, we sought to identify the B subunit responsible for targeting PP2A to DP. In an unbiased proteomics study, DP was identified as binding to the B subunit B55α^23^. Additionally, in both cell and mouse models, B55α loss resulted in impaired epidermal barrier development^24-27^. Based on these reports, we set out to address the extent to which B55α and DP associate in keratinocytes.

Toward this end, a co-immunoprecipitation (co-IP) assay was performed to pull down endogenous DP complexes using two different DP-directed antibodies targeting DP’s C- and N-termini. Both the C- and N-terminus targeting antibodies pulled down desmosomal cadherin Dsg3 as expected. In addition, B55α was also enriched in both IP complexes when compared to the IgG control (Figure 2A). A co-IP using an antibody targeting endogenous B55α similarly resulted in an enrichment in DP compared to control, but interestingly did not enrich for Dsg3 suggesting B55α may preferentially interact with DP (Figure 2B). To look at potential B55α-DP co-localization, immunofluorescence was performed on SCC9 cells co-staining B55α with either DP or other membrane-associated proteins including the desmosomal and adherens junction-associated armadillo protein PG, and the adherens junction cytoskeletal linker protein α-catenin. Staining revealed B55α more closely resembled DP’s localization patterns than PG or α-Catenin (Figure 2C). Co-localization was quantified using an object-based analysis tool, revealing significantly higher colocalization between B55α/DP than B55α/α-catenin (Figure 2D)^28^. Staining SCC9s transfected with B55α targeted siRNA confirmed that the membrane localized B55α signal is specific to B55α protein (Figure 2E). Notably, B55α loss was associated with a decrease in total amount of DP at the cell membrane suggesting B55α may regulate DP dynamics (Figure 2E). Additionally, loss of DP coincided with loss of membrane associated B55α, supporting the notion that B55α is physically anchored to DP at the membrane. To validate this interaction in cells, a proximity ligation assay (PLA) using antibodies targeting B55α and DP was performed in SCC9s transfected with siRNA targeting either a Scramble control, B55α, or DP^29^. Consistent with the colocalization analysis, the PLA produced a high amount of signal in the Scramble control cells that was lost with the knockdown of either B55α or DP, confirming that the PLA signal is due to close proximity between the two proteins within cells (Figure 2F).

**Figure 2:**
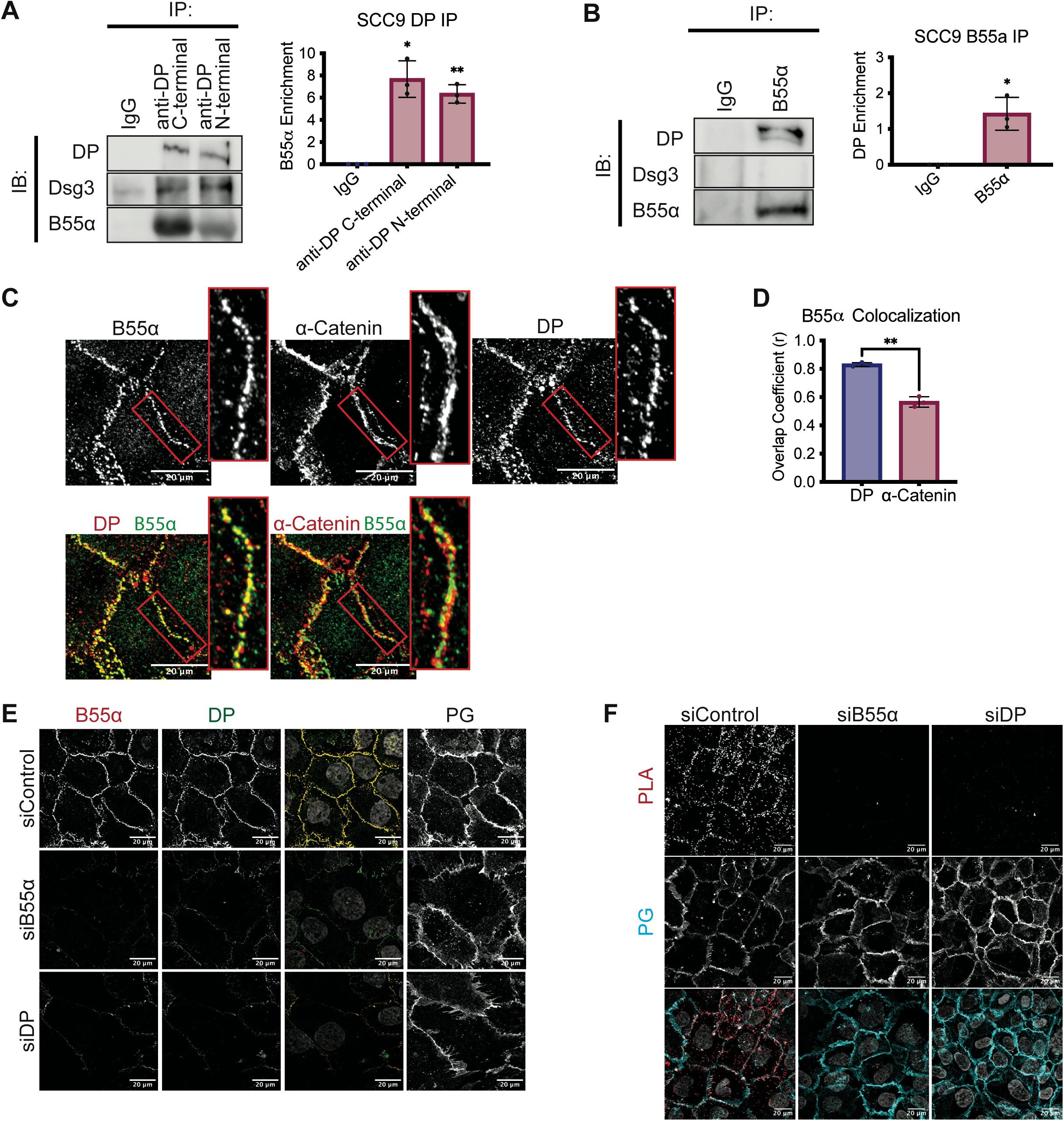
DP is found in complex with the PP2A regulatory subunit B55α in SCC9 cells. [A] Immunoprecipitation of endogenous DP using antibodies targeting DP’s C-terminus or N-terminus and blotting back for Dsg3 or B55α. [B] Immunoprecipitation of endogenous B55α and blotting back for DP or Dsg3. [C] SCC9 immunofluorescence co-stained for B55α, DP, and α-Catenin. Overlayed images are shown below. [D] Colocalization analysis of [C] as determined by an object-based colocalization analysis tool represented as overlap coefficient measurements. [E] SCC9s transfected with siRNA targeting B55α or DP and analyzed by immunofluorescence. [F] Proximity ligation analysis performed on SCC9 cell transfected with siRNA targeting either a scramble control, B55α, or DP. A fluorescence-based PLA signal was measure on fixed coverslips incubated with B55α and DP targeting antibodies. Statistical analyses were performed using a One-way Anova with multiple comparisons. * < 0.05; ** < .01.

To determine if a B55α/DP complex also exists in tissues, a 3D organotypic reconstructed epidermal raft model grown for 6 days was co-stained for B55α and DP. Consistent with the cellular data, B55α was found localized at the membrane in a pattern more closely resembling DP than PG or α-Catenin (Figure 3A-B, Supplemental Figure 2A-B). Furthermore, object-based co-localization analysis revealed B55α had a higher overlap coefficient with DP than with either PG or α-Catenin (Figure 3C). Staining human skin samples also showed B55α at the membrane within the intact epidermis in a pattern consistent with in vitro results, suggesting this interaction is maintained in vivo (Figure 3D-E, Supplemental Figure 2C). Lastly, to more directly interrogate the B55α-DP interaction in vivo, PLA was performed on human skin samples using B55α and DP directed probes and compared to samples treated with B55α antibody or IgG alone. Consistent with the cellular data, PLA signal was robustly detected compared to controls, further supporting the conclusion that B55α is in complex with DP within the intact stratified epidermis (Figure 3F).

**Figure 3:**
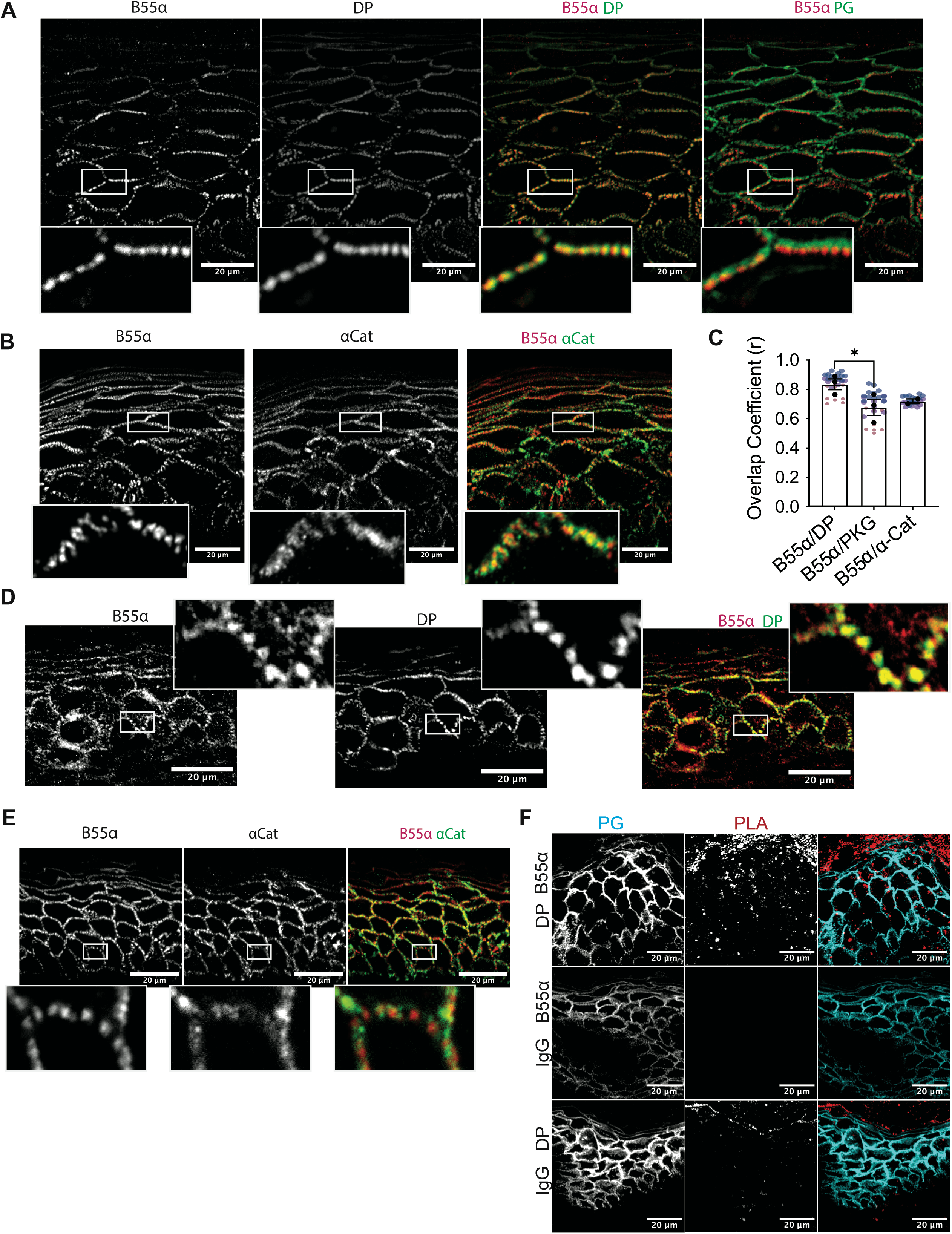
B55alpha is found localized to sites of intercellular membranes in the 3D epidermis in vitro and in vivo. [A-B] 3D organotypic reconstructed skin cultures grown for six days in culture and immunofluorescence staining of B55α, DP, and PG [A] and B55α, α-Catenin, and PG [B] was performed. [C] Colocalization analysis of [A-B] as determined by an object-based colocalization analysis tool represented as overlap coefficient measurements. B55α/PG colocalization measurements are from an n=3; B55α/α-catenin colocalization measurements are from an n=2. [D-E] Human skin samples were stained for B55α, DP, and PG [D] and B55α, α-Catenin, and PG [E] using immunofluorescence. [F] Proximity ligation analysis performed on fixed human skin samples. A fluorescence-based PLA signal was measure on fixed slides incubated with either B55α and DP targeting antibodies or B55α and IgG. Statistical analyses were performed using a One-way Anova with multiple comparisons. * < 0.05; ** < .01.

### PP2A-B55α regulates desmosome-mediated cell adhesion

Our group previously reported that ectopic expression of constitutively hypo-phosphorylated DP C-terminus increases adhesion strength^17^. Therefore, we were interested in identifying the role of PP2A-B55α in regulating keratinocyte adhesion through DP’s phospho-regulation. A dispase-based adhesion assay was performed on SCC9 cells in which either the PP2A catalytic subunit (PP2A-C) or the B55α regulatory subunit was knocked down using siRNA. Cells were grown for 5 days in fully confluent monolayers and dispase treatment was used to lift cell monolayers off the culture plate before transporting monolayers into tubes to undergo mechanical stress through rotation. Adhesion strength was determined by recording the level of fragmentation of the monolayers. Silencing of either PP2A-C or B55α resulted in significantly more monolayer fragments when compared to controls (Figure 4A-C). Notably, siDP treated cells resulted in complete fragmentation of cell monolayers suggesting PP2A-B55α loss only partially disrupts DP function. Collectively, these studies present PP2A-B55α as a molecular module capable of modifying the adhesive strength of keratinocytes in response to their changing mechanical environments.

**Figure 4:**
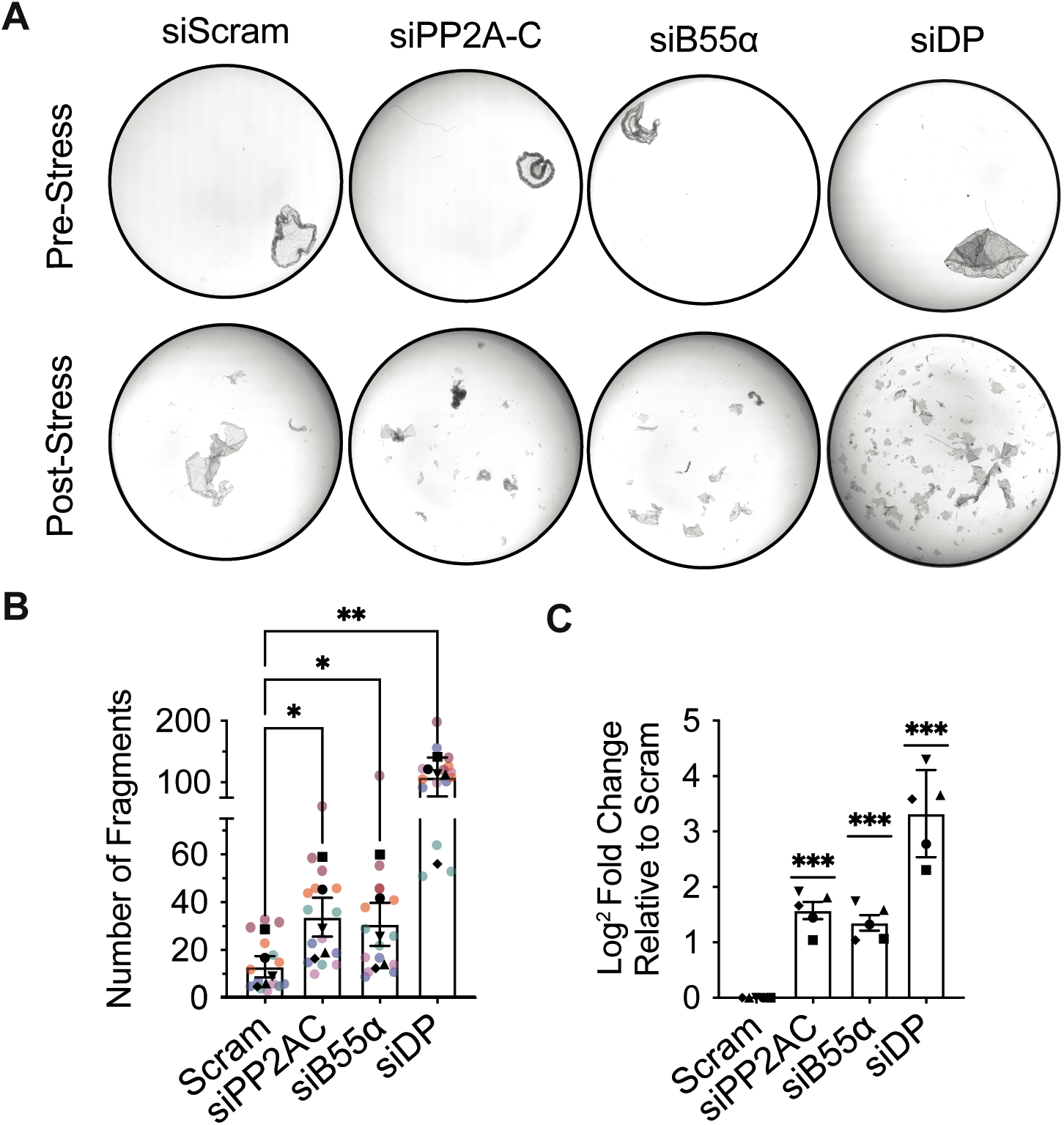
PP2A-B55alpha regulates epithelial cell adhesion and barrier through DP. [A] A dispase-adhesion assay was performed in SCC9s transfected with siRNA targeting scramble, B55α, or DP and grown in normal HCM for 5 days. Images represent SCC9 monolayers pre-(left) and post-(right) exposure to mechanical stress. Monolayer fragmentation from [A] was quantified and represented as total fragment number [B] or as a Log2 fold change normalized to their experimental scramble control [C]. Statistical analyses were performed using a One-way Anova with multiple comparisons from n=5. * < 0.05; ** < .01.

## DISCUSSION

By identifying the phosphatase that acts on the phospho-motif at the DP C-terminus we have filled a major gap in our understanding of how the desmosome-intermediate filament connection is tuned through regulation of DP phosphorylation state. Our work shows that the PP2A-B55α holoenzyme is associated with the desmosomal cytoskeletal linker DP and loss of its activity results in accumulation of hyper-phosphorylated DP concentrated at cell-cell junctions. We showed further that DP phosphorylation induced by inhibition of PP2A-B55α weakens intercellular adhesion, presumably through a decrease in DP-IF connection strength. We previously demonstrated that the enzymes GSK3β and PRMT-1 cooperate to mediate DP phosphorylation^14^. Together with the current work, we propose an updated regulatory mechanism whereby the PP2A-B55α and PRMT-1/GSK3β form a molecular switch through the phospho-regulation of DP’s C-terminus to fine-tune the DP-IF connection that controls cell adhesive and tensile strength (Figure 5).

**Figure 5:**
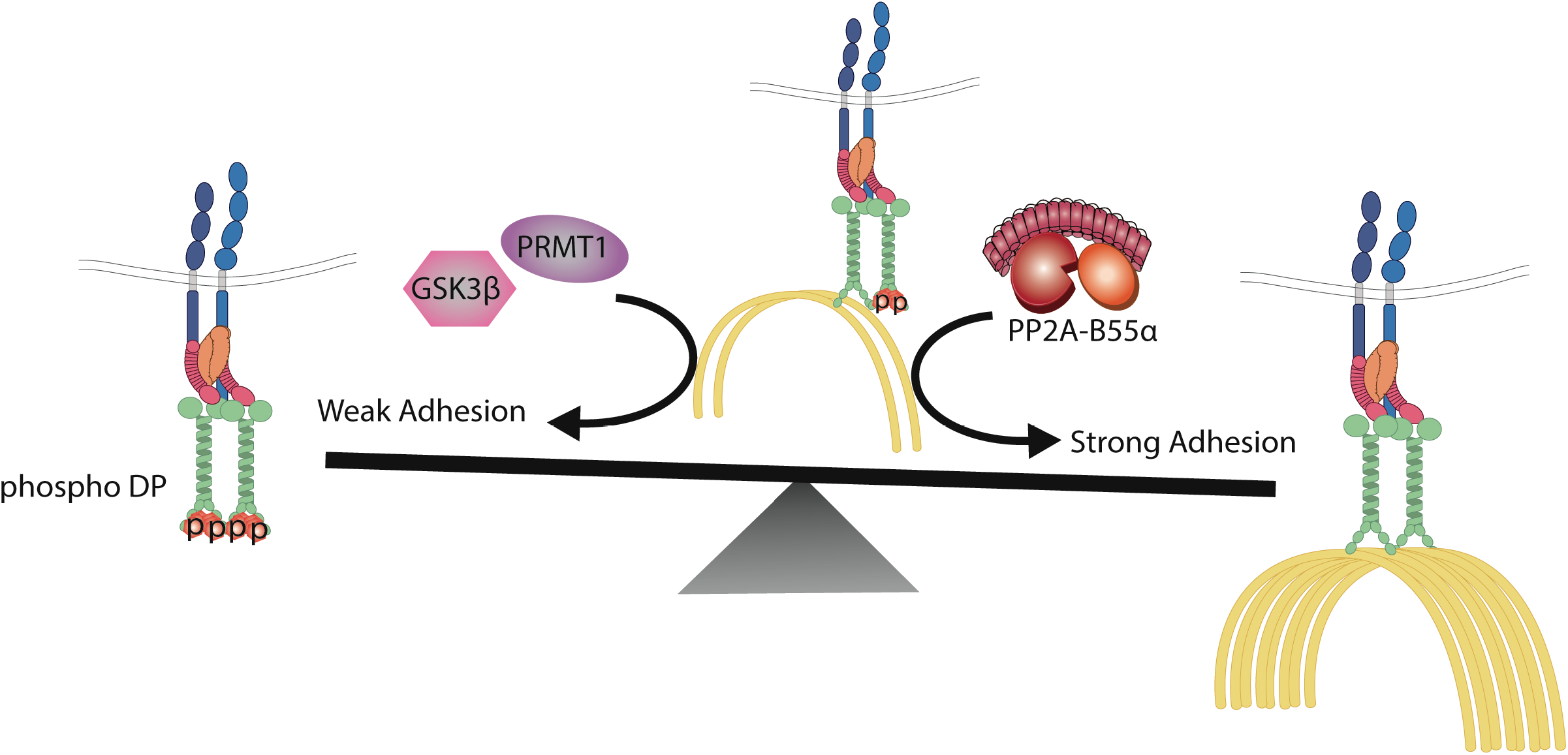
A balance between PP2A-B55alpha and GSK3β/PRMT1 activities on DP’s C-terminus controls desmosome-adhesion strength through regulating the DP-IF connection. A schematic representing the regulatory switch controlling desmosome-mediated adhesion through the phospho-regulation of the DP-IF interaction. Albrecht et al. previously have previously identified GSK3β and PRMT1 as cooperating to induce the phosphorylation of DP’s C-terminus that inhibits DP-IF binding and weakens desmosomal adhesion^14^. Our work has uncovered of PP2A-B55α as an opposing regulatory node capable of dephosphorylating DP’s C-terminus, inducing increased cell adhesion potentially through in increased association between DP and IF.

PP2A-B55α activity has been associated with regulating several cellular processes including cell growth, DNA replication, mitotic exit, cytokinesis, and microtubule integrity^30,31^. Its most well characterized substrates include the tubulin binding protein tau, the spindle regulatory protein PRC1, the pocket protein p107, and many of the CDK1 cell cycle substrates. With the strong overlap of B55α substrates and cell cycle regulation, PP2A-B55α has primarily been thought of as a cell cycle regulatory enzyme. Our work presents a potential role for the cell cycle regulator PP2A-B55α in non-replicating differentiated keratinocytes that experience high levels of intercellular tension requiring a strong desmosome-IF anchor and no longer require high levels of cell cycle regulation.

Recent work in cardiomyocytes and simple epithelia revealed desmosomes to be mechano-sensitive structures, which respond to mechanical stress by recruiting desmosome proteins to sites of cell-cell adhesion^32,33^. Despite these reports, it is unclear how the desmosome may differentially impact epidermal mechanics across the stratified epidermis. One potential mechanism could be through its cytoskeletal connections. Consistent with this idea, interference with the DP-IF interaction inhibits tight junction formation in 3D models, supporting its importance for epidermal barrier function^34,35^. Given our previous observation that a hypo-phosphomimic of DP increases epithelial sheet stiffness^19^, our data raise the possibility that tuning DP-IF interactions through PP2A-B55α mediated de-phosphorylation could contribute to the stiffness gradient recently reported to exist in the stratified epidermis^36^. In turn, increased tension in the superficial epidermis would help maintain tight junctions in the superficial stratum granulosum 2 (SG2). This is consistent with previous studies showing PP2A catalytic activity and the B55α regulatory subunit specifically are required for normal epidermal development, barrier function and specifically tight junction formation in cellular and in vivo models of PP2A loss^24-27^. Furthermore, pharmacologically-activation of PP2A improves epidermal barrier function when applied topically^37^. Together, these data uncover a possible mechanism to explain how the PP2A-B55α holoenzyme regulates both the epidermal barrier and tissue mechanics.

The importance of the DP-IF association is highlighted by the severe cardio-cutaneous diseases associated with mutations in DP’s C-terminus domain that interfere with the IF binding site and phospho-regulatory motif^9,13^. These include the cardiocutaneous disease Carvajal syndrome, associated with a truncation mutation at DP’s C-terminus, and the desmosome-related disease AC^10,38,39^. AC is associated with mutations in several desmosome components including a point mutation within DP’s C-terminus that interferes with the proper regulation of the phospho-regulatory motif that we have identified as a target of PP2A-B55α. Furthermore, ectopic expression of a constitutively hypo-phosphorylated DP mutant was sufficient to restore IF distribution and preserved cell-cell adhesion in a desmosome deficiency disease model of pemphigus vulgaris^40^. Notably, only half of all AC cases have been linked to mutations in known desmosome components^39^. The identification of PP2A-B55α as a regulator of DP’s C-terminus phospho-motif raises the possibility that mutations in the PP2A-B55α complex could be a candidate driving desmosomal diseases in patients without an identified desmosome mutation. Finally, previously published collaborative work showing PP2A inhibition increased phosphorylation of a conserved sequence in the more ubiquitously expressed plakin family relative Plectin, raises the possibility that B55α could play a broader regulatory role for plakins in different tissue types including cardiac and skeletal muscle^41^.

In summary, we have identified PP2A-B55α as a newly recognized component of a molecular switch also containing PRMT-1/GSK3β. Through its ability to negatively regulate a critical phospho-motif on DP’s C-terminus we propose that PP2A-B55α tunes desmosome-mediated adhesion to provide desmosomes with dynamic properties necessary for the development and maintenance of the epidermal barrier.

## MATERIAL AND METHODS

### Cell lines, culture conditions, and treatments

Human-derived oral squamous cell carcinoma SCC9 cells (a gift from J. Rheinwald, Harvard Medical School, Boston, MA) were cultured in DMEM/F12, 10% FBS, and 1% penicillin/streptomycin. Immortalized human keratinocyte HaCaT cells were cultured in DMEM, 10% FBS, and 1% penicillin/streptomycin. Cell lines were maintained at 37°C in a humidified atmosphere of 5% CO_2_. NHEKs are isolated from neonatal foreskin provided by the Northwestern University Skin Biology and Disease Resource-based Center (SBDRC) as previously described ^42^. NHEK’s were maintained in M154 (Thermo Fisher Scientific) growth media supplemented with 0.07 mM CaCl_2_, human keratinocyte growth supplement (HKGS; Thermo Fisher Scientific), gentamicin and amphotericin B. NHEKs were used to generate 3D reconstructed epidermal organotypic cultures as previous described ^42^.

For siRNA transfection, cells were plated and grown overnight to 60% confluency before transfection with Dharmafect (Thermo Fisher Scientific). For drug treatment studies, cells were grown at confluency for 3 days before treatment with OA (50nM or 100nM), Tautomycin (250nM), or DMSO for 3 hours. NHEKs were grown to confluency and incubated in growth media supplement with 1.2 mM CaCl_2_ for 2 days prior to drug treatment.

### Immunofluorescence and microscopy

Cells plated on glass coverslips were fixed either in anhydrous ice-cold methanol for 3 min on ice to visualize phospho S2849 DP/total DP staining or 4% paraformaldehyde (PFA) solution for 20 min at room temperature followed by anhydrous ice-cold methanol for 3 min on ice to visualize B55α. Cells were blocked with either 5% goat serum or 1% BSA and 1% donkey serum. All samples were mounted onto glass slides with ProLong Gold antifade reagent (Thermo Fisher Scientific). 3D reconstructed skin and human skin samples were stained from frozen embedded samples and fixed as described above. For PLA analysis, samples were fixed as described above and PLA was performed as described in Hegazy et. al^29^.

Apotome images were acquired using ZEN 2.3 software with an epifluorescence microscope system (Axio Imager Z2, Carl Zeiss) fitted with an X-Cite 120 LED Boost System, an Apotome.2 slide module, Axiocam 503 Mono digital camera, and a Plan-Apochromat 40x/1.4, Plan-Apochromat 63x/1.4 objective, Plan-Apochromat 100x/1.4 objective (Carl Zeiss). Images are processed using ImageJ software. Colocalization analysis was performed using the JaCoP ImageJ plugin^28^.

### Western blot analysis

Whole cell lysates were generated using Urea Sample Buffer (8 M urea, 1% SDS, 60 mM Tris, pH 6.8, 5% b-mercaptoethanol, 10% glycerol). Proteins were separated by SDS-PAGE electrophoresis and transferred to nitrocellulose membranes. 5% milk was used to block membranes and dilute primary and secondary antibodies. Immunoreactive proteins were visualized using chemiluminescence or LiCOR fluorescence secondary antibodies.

Phosphate-affinity SDS-PAGE was performed using Phos-tag (Wako Pure Chemical Industries). The procedure was performed following the manufacturer’s protocol (http://www.phos-tag.com). 50 μM of phos-tag acrylamide peptide is added during the preparation of a 15% wt/vol polyacrylamide gel. Gel electrophoresis was run at 15mAmps for 5.5 hours and transferred overnight onto a PDVF membrane. Subsequent immunoblotting was performed as described above. Quantification of bands was done using Image Lab software (Bio-Rad Laboratories).

### Antibodies and reagents

The following primary antibodies were used: NW6 Rabbit anti-DP C-terminal(Green Lab^43^); NW161 Rabbit anti-DP N-Terminal (Green Lab^44^); 115F Mouse anti-DP C-terminal (Sigma, Gift from D. Garrod^45^); anti–phospho-S2849 DP (Green Lab^46^); anti-Desmoglein 3 5G11(Sigma-Aldrich); 1407 Chicken anti-PG (Aves Laboratories); 2G9 Mouse anti-B55α (Cell Signaling Technology); PA5-18512 Goat anti-α-Catenin (Thermo Fisher Scientific); PA5-17443 Rabbit anti-α-Catenin (Thermo Fisher Scientific); mouse anti-S-Tag (EMD Millipore); Mouse anti-GAPDH (Santa Cruz Biotechnology); 12G10 Mouse anti-α-tubulin (Developmental Hybridoma Studies Bank). The following secondary antibodies were used: Goat anti-Mouse IgG HRP (Cell Signaling Technologies); Goat anti-Rabbit IgG HRP (Cell Signaling Technology); Goat anti-Mouse conjugated with Alexa Fluor-488 (Thermo Fisher Scientific); Goat anti-Mouse conjugated with Alexa Fluor-568 (Thermo Fisher Scientific); Goat anti-Chicken conjugated with Alexa Fluor-647 (Thermo Fisher Scientific); Donkey anti-Goat conjugated with Alexa Fluor-488 (Thermo Fisher Scientific); Donkey anti-Mouse conjugated with Alexa Fluor-568 (Thermo Fisher Scientific); Donkey anti-Rabbit conjugated with Alexa Fluor-647 (Thermo Fisher Scientific).

### Co-IP Assay

Cells grown at confluency for 3 days were lysed in NP40 buffer on ice for 30 minutes. Supernatant was collected by centrifugation and incubated overnight at 4°C rotating with anti-DP antibody or anti-B55α antibody conjugated to agarose (Sant Cruz Biotechnology). Agarose conjugated Protein A/G was rotated for 1 hour at 4°C before centrifugation and washes in 1x NP40 buffer. Samples were resuspended in 3x Lamelli Buffer and boiled at 100°C for 10 min before western blot analysis was performed.

### Dispase Assay

Cell monolayers were grown at confluency for 5 days in triplicate in 12-well plates. Cells were washed in PBS and treated with 2.4 U/mL of dispase (Sigma Millipore) diluted in PBS containing Ca^2+^ for 30 min. Lifted monolayers were placed in 15mL conical tubes containing PBS and subjected to mechanical stress by mechanical rotation. Resulting fragments were returned to 12-well dishes and images using a dissecting microscope (MZ6; Leica).

## ACKNOWLEDGEMENTS

We thank members of the Green lab for their valuable feedback and discussion. Research reported in this publication was supported by Northwestern University Skin Biology & Diseases Resource-Based Center of the National Institutes of Health under award number P30AR075049. Imaging work was performed at the Northwestern University Center for Advanced Microscopy generously supported by NCI CCSG P30 CA060553 awarded to the Robert H. Lurie Comprehensive Cancer Center. Northwestern University’s Pathology Core Facility (PCF) and Mouse Histology and Phenotyping Laboratory (MHPL) performed the sectioning of human skin samples and reconstructed 3D skin equivalents. This work was supported by NIH R01 AR43380, NIAMS R01 AR041836, and NCI R01 CA228196 to KJG. A.L.P was supported by T32AR060710.

## FIGURE LEGENDS

**Supplemental Figure 1**

HaCaT’s [A] and NHEKs [B] were treated with either DMSO, 50nM OA or 250nM Tautomycin for 3 hours. Expression levels of S2849 phosphorylated and total DP were analyzed by immunofluorescence staining. Staining intensity was quantified at the membrane using PG stain as a mask. Statistical analyses were performed on normalized data using a One sample t comparison to 1. * < 0.05; ** < .01.

**Supplemental Figure 2**

[A] Staining intensities of B55α, DP, and PG across the 3D-reconstructed skin is shown using a HEAT map representation. [B] PG, B55α, and α-catenin staining in 3D reconstructed skin from Figure 3B. [C] PG, B55α, and α-catenin staining in human skin samples from Figure 3D.

