## Supplementary figures and images for "PP2A-B55α controls keratinocyte adhesion through dephosphorylation of the Desmoplakin C-terminus"

### Supplemental Figure 1

#### Supplemental Figure 1

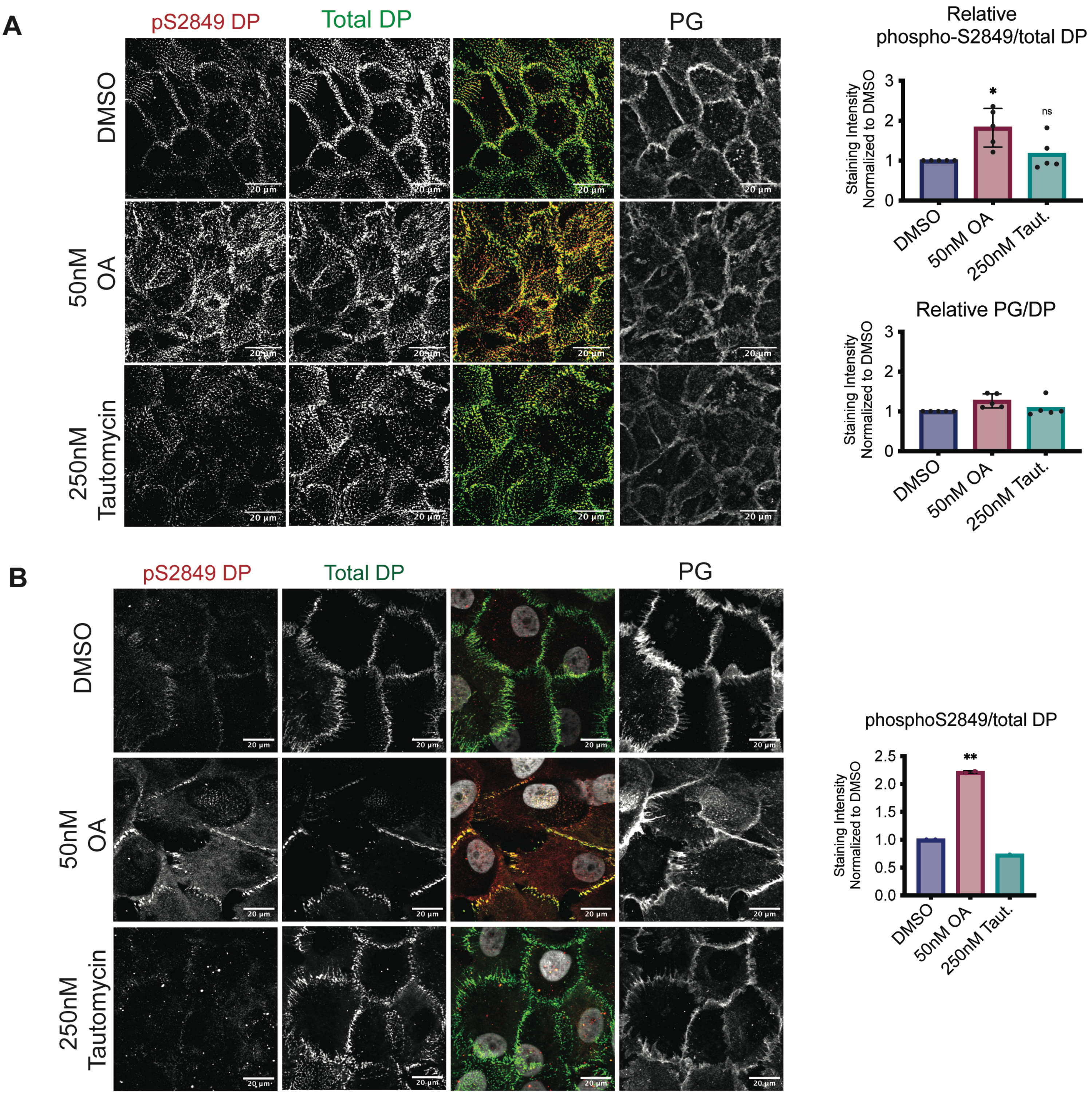

### Supplemental Figure 2

#### Supplemental Figure 2

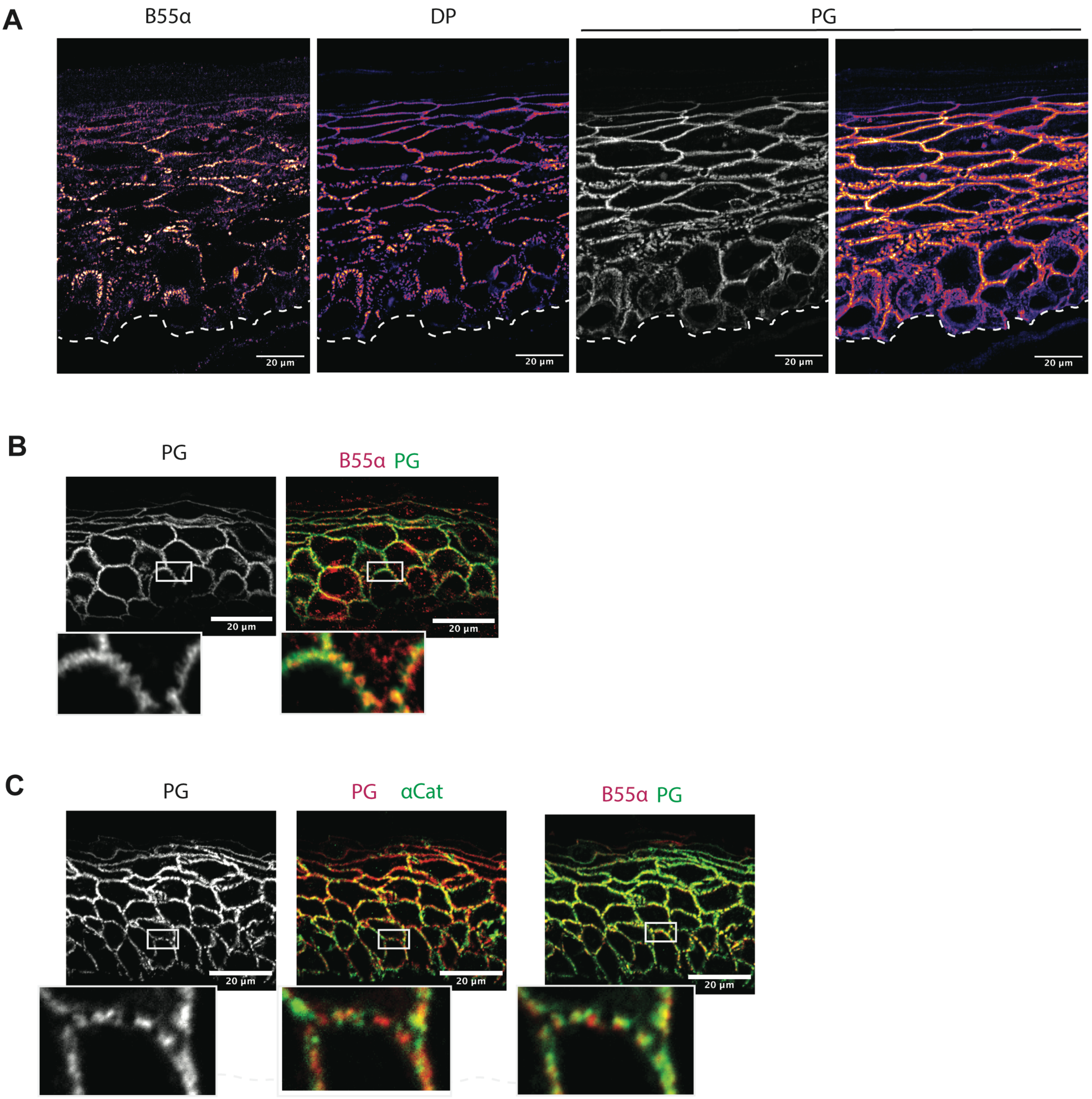
